# Rose Extract Treatment on the CD4+ T lymphocytes

**DOI:** 10.1101/2021.04.06.438604

**Authors:** Mark Christopher Arokiaraj, Eric Menesson

## Abstract

**Background and Aims:** To study the effect of rose extract on CD4+T lymphocytes, and assess the cytokines response after cell treatment. In our previous study on endothelial cells, the rose extract reduced the secretion of inflammatory markers significantly.

**Methods:** The red rose extract used in this study was prepared and stored until use at -20°C. T cells were seeded in 96-well plates at 313500 cells/well in 100μl of cell culture medium in duplicate, one half of the wells were used for biomarkers screening in the culture medium, and the other half for cytotoxicity assay. 24h after plating, the cells were treated in duplicate with 100μl of red rose extract diluted at 0.5%, 0.1%, 0.05%, 0.01% and 0.005% (v/v) in cell culture medium or with culture medium only as control for 72h. Some other wells were for untreated cells, and cells treated with rose extract at 0.005% for 48h incubation time. After 48h and 72h, the corresponding wells were used for the cytotoxicity assay and from the duplicate wells, the cell culture media were collected and stored at -80°C until the biomarkers screening assay.

**Results:** Cytotoxicity assay revealed insignificant changes. IFN-gamma, MCP-1, GRO, RANTES and TIMP, Angiopoietin 1 and MMP-9 were elevated. Except MMP-9 which had fold changes >2 other cytokines were minimally elevated at various concentrations and timing of rose extract treatment. None of the cytokines were less than 0.8-fold.

**Conclusions:** Unlike in the endothelial cells, there is mild elevation in few inflammatory markers on T lymphocytes treatment by rose extract. Further studies need to be performed to estimate the clinical relevance.

## Introduction

Immune diseases are common in clinical practice. Most medical disorders have some immunological background. Hence, positive immune modulation with reduction of inflammation has potential advantages as a therapy. The current immune modulators reduce lymphocyte proliferation and function, which can be deleterious. The study was performed in search of a novel immune-modulator, which can potentially reduce the risk of reduction in T-cell lymphocyte function. In our previous studies, we showed the anti-inflammatory effect of the rose extract on the endothelial cells *in-vitro*.^1^ It also had effects of inhibition of SARS-CoV-2 spike protein on target ACE-2 receptors.^2^ Currently existing most anti-inflammatory agents like steroids,^3^ tacrolimus,^4^ MMF^5,6^ etc., reduce lymphocyte cell proliferation, and decrease inflammation. Monoclonal antibodies like rituximab,^7-9^ infliximab^10^ etc., are also associated with various side effects. Rituximab, which is widely used in various lymphoproliferative disorders and autoimmune disorders has anti-CD20 and also increases IL10.

## Methods

### HUVEC Treatment

The red rose extract used in this study was prepared for our previous study ^1,11^ and stored until use at -20°C. Human peripheral blood CD4+ T cells (Cell Applications, ref. 6902-50a, lot 3298) were seeded in 96-well plates at 313500 cells/well (950000 cells/cm2) in 100μl of cell culture medium (Cell Applications, ref. 615-250) in duplicate, one half of wells used for biomarkers screening in the culture medium and the other half for cytotoxicity assay. 24h after plating, the cells were treated in duplicate with 100μl of red rose extract diluted at 0.5%, 0.1%, 0.05%, 0.01% and 0.005% (v/v) in cell culture medium or with culture medium only as control for 72h. Some other wells were for untreated cells and cells treated with rose extract at 0.005% for 48h incubation time. After 48h and 72h, the corresponding wells were used for the cytotoxicity assay and from the duplicate wells, the cell culture media were collected and stored at -80°C until the biomarkers screening assay. The biomarkers screening assay was performed with human angiogenesis antibody array G-series from Raybiotech, ref. AAH-ANG-G1 and AAH-ANG-G2.

### Cytotoxicity assay

After 48h incubation time, cell viability was assessed using « Cell Counting Kit–8 » (Dojindo EU Gmbh, ref. CK04). This kit uses WST-8 (2-(2-methoxy-4-nitrophenyl)-3-(4-nitrophenyl)-5-(2,4-disulfophenyl)-2H-tetrazolium, monosodium salt) which produces a water-soluble formazan dye upon bio-reduction in the presence of an electron carrier, 1-Methoxy PMS. WST-8 (10% of the medium volume in wells) was added to the media for 4h at 37°C. During the incubation time, it is bio-reduced by cellular dehydrogenases to an orange formazan product that is soluble in culture medium. Then, the amount of formazan produced is directly proportional to the number of living cells. The absorbance of formazan was measured at 450 nm and enables the calculation of viable cells percentage for each treatment compared to non-treated cells (Table 1).

### Profiling of secreted cytokines

The technique of quantibody assay is similar to the method described in our previous studies.^1,11^ During the incubation time, the volume of medium varied differently in each well and the volume collected might be reduced. In order to normalize the profiling results, cell culture medium was added to the different collected media to reach 200μl before performing the profiling assay. Then, the medium samples were tested undiluted on arrays.

## Results of profiling

### Cytotoxicity assay

The results of cytotoxicity assay showed there was no strong effect of treatment on the viability of the T cells even though the viability went down until 87% for cells treated 72h with 0.01% rose extract. Indeed, such a decrease of viability in one tested replicate is not statistically representative of a cytotoxicity effect of the rose extract. Figure 1 shows the results of cytotoxicity assay.

**Figure 1:**
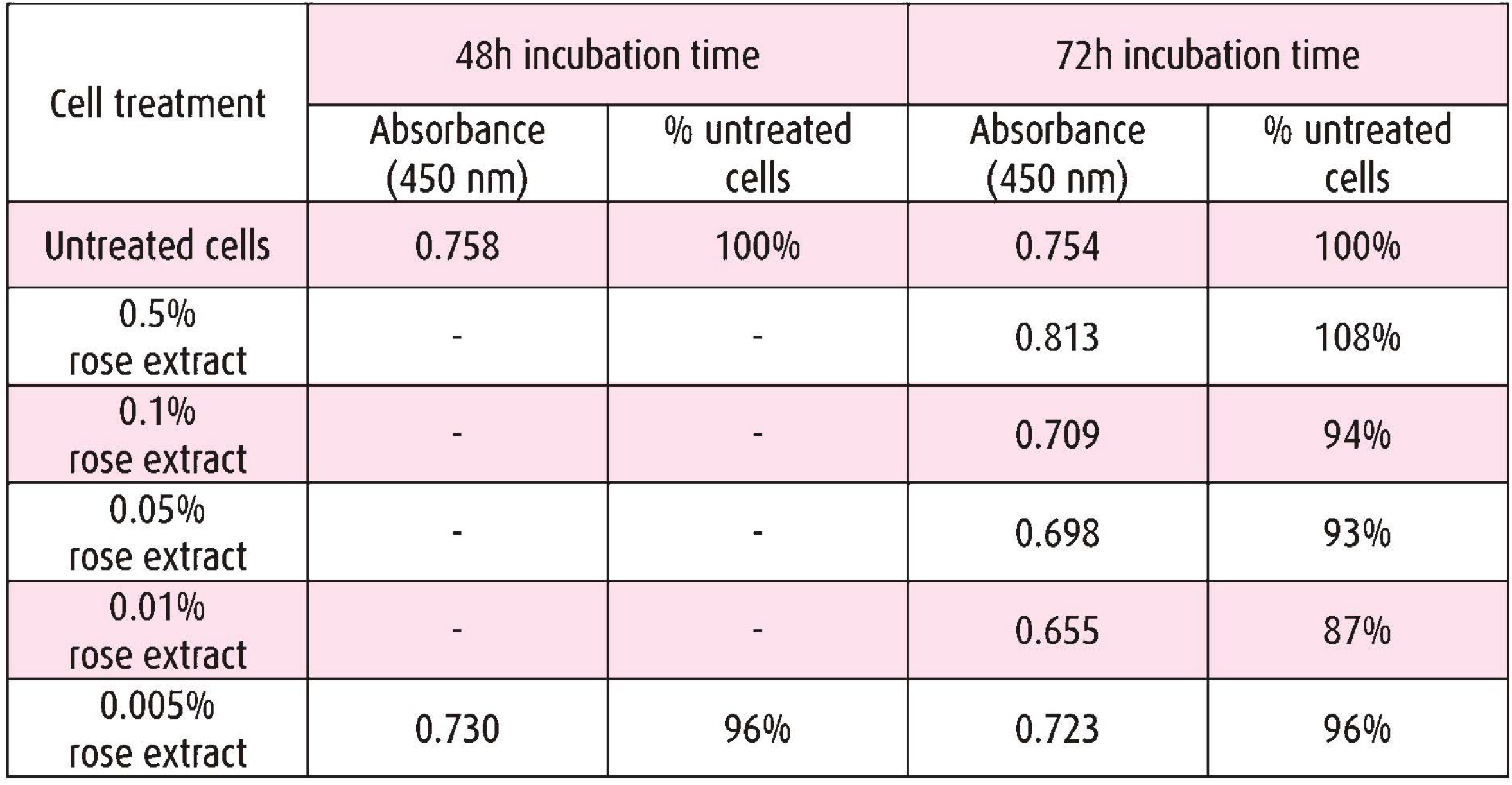
The results of cytotoxicity assay based on absorbance observed due to orange formazan product.

### Cytokines

IFN-gamma, MCP-1, GRO, RANTES and TIMP, Angiopoietin 1 and MMP-9 were elevated. Except MMP-9 which had fold changes >2, other cytokines were mildly elevated at various concentrations and timing of rose extract treatment (Table 2 and 3). GRO was mainly elevated at higher (0.5%) and lower (0.005%) concentrations. MCP-1 and RANTES showed an increasing trend with decreasing levels of rose-extract concentrations. IL-6 showed an higher values in higher (0.5%) and lower dilutions (0.005%). The results also showed that the cytokines IL-6, IL-10, IL-1a, TGF-b1, TIMP-2, Angiopoietin-2, G-CSF, and VEGF R2 were enough detected to enable the calculation of one-fold change at least.

**Table 2:** Results of secreted cytokines profiling: fold changes of treated cells vs untreated cells at both incubation times for AAH-ANG-G1 microarray. Target not detected (-)

|  | 0,005% 48h<br>vs<br>Untreated cells 48h | 0,5% 72h<br>vs<br>Untreated cells 72h | 0,1% 72h<br>vs<br>Untreated cells 72h | 0,05% 72h<br>vs<br>Untreated cells 72h | 0,01% 72h<br>vs<br>Untreated cells 72h | 0,005% 72h<br>vs<br>Untreated cells 72h |
| --- | --- | --- | --- | --- | --- | --- |
| Angiogenin | - | - | - | - | - | - |
| EGF | - | - | - | - | - | - |
| ENA-78 | - | - | - | - | - | - |
| b FGF | - | - | - | - | - | - |
| GRO | 0.92 | 1.84 | 1.17 | 1.05 | 1.27 | 1.61 |
| IFN-gamma | 0.96 | 1.19 | 1.25 | 1.23 | 1.15 | 1.21 |
| IGF-I | - | - | - | - | - | - |
| IL-6 | 0.96 | 1.55 | 1.12 | 1.12 | 1.13 | 1.21 |
| IL-8 | - | - | - | - | - | - |
| LEPTIN | - | - | - | - | - | - |
| MCP-1 | 1.12 | 1.08 | 1.33 | 1.08 | 1.39 | 1.75 |
| PDGF-BB | - | - | - | - | - | - |
| PIGF | - | - | - | - | - | - |
| RANTES | 0.92 |  | 0.80 | 1.14 | 1.45 | 1.63 |
| TGF-beta1 | 0.93 | 1.05 | 1.07 | 0.97 | 1.01 | 1.13 |
| TIMP-1 | 0.84 | 1.04 | 1.24 | 1.04 | 1.11 | 1.23 |
| TIMP-2 | - | - | 1.20 | - | 1.05 | 1.16 |
| Thrombopoietin | - | - | - | - | - | - |
| VEGF | - | - | - | - | - | - |
| VEGF-D | - | - | - | - | - | - |

**Table 3:** Results of secreted cytokines profiling: fold changes of treated cells vs untreated cells at both incubation times for AAH-ANG-G2 microarray. Target not detected (-).

|  | 0,005% 48h<br>vs<br>Untreated cells 48h | 0,5% 72h<br>vs<br>Untreated cells 72h | 0,1% 72h<br>vs<br>Untreated cells 72h | 0,05% 72h<br>vs<br>Untreated cells 72h | 0,01% 72h<br>vs<br>Untreated cells 72h | 0,005% 72h<br>vs<br>Untreated cells 72h |
| --- | --- | --- | --- | --- | --- | --- |
| Angiopoietin-1 | 0.72 | 0.69 | 1.30 | 1.29 | 1.35 | 1.18 |
| Angiopoietin-2 | 0.68 | 0.94 | 1.01 | 1.05 | 0.92 | 0.79 |
| Angiostatin | - | - | - | - | - | - |
| Endostatin | - | - | - | - | - | - |
| G-CSF | 1.04 | - | - | - | - | - |
| GM-CSF | - | - | - | - | - | - |
| I-309 | - | - | - | - | - | - |
| IL-10 | 1.14 | - | - | - | - | 0.94 |
| IL-1alpha | 1.05 | 1.17 | 1.09 | 1.00 | 0.96 | 0.93 |
| IL-1beta | - | - | - | - | - | - |
| IL-2 | - | - | - | - | - | - |
| IL-4 | - | - | - | - | - | - |
| I-TAC | - | - | - | - | - | - |
| MCP-3 | - | - | - | - | - | - |
| MCP-4 | - | - | - | - | - | - |
| MMP-1 | - | - | - | - | - | - |
| MMP-9 | - | 2.68 | 1.81 | 1.71 | 2.62 | 2.23 |
| PECAM-1 | - | - | - | - | - | - |
| Tie-2 | - | - | - | - | - | - |
| TNF-alpha | - | - | - | - | - | - |
| u PAR | - | - | - | - | - | - |
| VEGF R2 | 0.98 | 1.13 | 1.08 | 0.96 | 0.93 | 0.84 |
| VEGF R3 | - | - | - | - | - | - |

## Discussion

The results showed that the cytokines GRO, IFNg, IL-6, MCP-1, RANTES, TGF-b1, TIMP-1, TIMP-2, Angiopoietin-1, Angiopoietin-2, G-CSF, IL-10, IL-1a, MMP-9 and VEGF R2 were enough detected to enable the calculation of one-fold change at least. In this study, the CD4+ T cells have shown a mild stimulatory effect on the cells. Our previous results of rose extract treatment on the endothelial cells showed a reduction in the levels of cytokine levels and the inflammatory markers. In the clinical scenario it could be advantageous. A reduction in the inflammation of the endothelial cells is combined with stimulatory effect on the lymphocytes, which is uniquely observed in this study. The cytotoxicity assay in this study revealed insignificant changes with minimal reduction in the viability of cells. Absence of cytotoxicity of the rose extract even in higher concentrations has potential advantages to evolve as a therapy in the future.

### Elevated cytokines

IFN-gamma, MCP-1, GRO, RANTES and TIMP1, Angiopoietin 1 were elevated, and MMP-9 were elevated more than 2-fold. Interferon gamma^12,13^ is playing a key role in immune functions involving a broad spectrum of diseases. MCP-1^14^ and GRO^15^ are chemotactic cytokines especially for the mononuclear cells. RANTES^16^ and TIMP1^17^ are inflammatory mediators in acute and chronic inflammation. Angiopoietin 1 is involved in master regulation of angiogenesis.^18,19^ MMP9 is involved in vascular remodelling of angiogenesis, wound repair and neutrophil migration across basement membrane.^20,21^

### Minimally elevated cytokines

IL6,^22,23^ IL10 ^24,25^ and IL1a^26^ are major regulators in inflammation, immunity and infections. Angiopoietin 2^27^ and VEGF R2^28^ are involved in angiogenesis and inflammation. G-CSF has a role in granulopoiesis and anti-inflammation.^29^ TIMP-2 ^30,31^ and TGF-b are mildly elevated.^32^ TIMP2 actively attenuates the negative remodelling and TGFb2 is involved in tissue repair.^33^ The catalytic effects of the rose extract on the immune system also needs to be studied, and since the performance invitro is unique it could act as a ‘salt’ in the immune functions, though speculative in the current scenario.

### Limitations

This study was performed invitro with limited number of samples and numbers. Further extensive evaluation of the extract needs to be performed in large numbers *invitro* and later in animal models. The side-effect profile of the extract also needs to be evaluated. Also, the feedback mechanism, which could exist in these T-cells in immune regulation need to be studied in vivo.^34.^ The quality of the rose extract needs to be standardised by HPLC etc.

### Conclusions

In this study, the effect of red rose extract on the secretion of cytokines from T cells CD4+ was assessed. The secreted cytokines were profiled after 48h or 72h of incubation. The results showed that the cytokines GRO, IFNg, IL-6, MCP-1, RANTES, TGF-b1, TIMP-1, TIMP-2, Angiopoietin-1, Angiopoietin-2, G-CSF, IL-10, IL-1a, MMP-9 and VEGF R2 were enough detected to enable the calculation of one-fold change at least. There was no significant fold change over 2 or lower 0.5 except MMP-9 with fold change often over 2. However, a significant fold change variation between different concentrations of rose extract and untreated cells should be also considered like RANTES or to a lesser extent MCP-1, IL-1a, VEGF R2.

## Author contributions

MCA conceived the idea and method, designed the study, interpreted results and wrote the paper. EM prepared the methods protocol, performed the experiments and derived the results.

## Bibliography

1. Arokiaraj MC, Menesson E. Novel anti-inflammatory and immunomodulation effects of Rose on the endothelium in normal and hypoxic invitro conditions. Angiologia e Cirurgia Vascular. 2019;15(4):238–248.

2. Arokiaraj, Mark Christopher, & Menesson, Eric. (2020). Rose and in-vitro Inhibition of Sars-CoV-2 spike: ACE-2 interaction (Version 1). http://doi.org/10.5281/zenodo.3886767.

3. Creed T, Lee R, Newcomb P, di Mambro A, Raju M, Dayan C. The Effects of Cytokines on Suppression of Lymphocyte Proliferation by Dexamethasone. The Journal of Immunology. 2009;183(1):164–171.

4. Härtel C, Schumacher N, Fricke L, Ebel B, Kirchner H, Müller-Steinhardt M. Sensitivity of whole-blood T lymphocytes in individual patients to tacrolimus (FK 506): impact of interleukin-2 mRNA expression as surrogate measure of immunosuppressive effect. Clin Chem. 2004 Jan;50(1):141–51. doi: 10.1373/clinchem.2003.024950. PMID: 14709642.

5. Ritter ML, Pirofski L. Mycophenolate mofetil: effects on cellular immune subsets, infectious complications, and antimicrobial activity. Transpl Infect Dis. 2009;11(4):290–297. doi:10.1111/j.1399-3062.2009.00407.x

6. Guzera M, Szulc-Dabrowska L, Cywinska A, Archer J, Winnicka A (2016) In Vitro Influence of Mycophenolic Acid on Selected Parameters of Stimulated Peripheral Canine Lymphocytes. PLoS ONE 11(5): e0154429. https://doi.org/10.1371/journal.pone.0154429

7. Weiner GJ. Rituximab: mechanism of action. Semin Hematol. 2010;47(2):115–123. doi:10.1053/j.seminhematol.2010.01.011

8. In addition to B-cell depletion, rituximab can modulate the immune response by inducing cytokine secretion, especially IL-10 and MIP-1β. Rituximab-induced MIP-1β secretion depends on the combined presence of B cells and FcR-bearing cells, especially NK cells.

9. Kamburova EG, van den Hoogen MW, Koenen HJ, Baas MC, Hilbrands LB, Joosten I. Cytokine Release After Treatment With Rituximab in Renal Transplant Recipients. Transplantation. 2015 Sep;99(9):1907–11. doi: 10.1097/TP.0000000000000515. PMID: 25675201.

10. Krupa L, Kennedy H, Jamieson C, Fisher N, Hart A. The Reasons for Discontinuation of Infliximab Treatment in Patients with Crohn’s Disease: A Review of Practice at NHS Teaching Hospital. ISRN Gastroenterology. 2011;2011:1–3. DOI: 10.5402/2011/672017

11. Arokiaraj MC, Menesson E, Feltin N. Magnetic iodixanol -a novel contrast agent and its early characterization. J Med Vasc. 2018 Feb;43(1):10–19. doi: 10.1016/j.jdmv.2017.11.002. Epub 2017 Dec 18. PMID: 29425536.

12. Zhao Zha, Felicitas Bucher, Anahita Nejatfard, Tianqing Zheng, HongkaiZhang, Kyungmo o Yea, Richard A. Lerner. Interferon-γ functions as a master switch. Interferon-γ functions as a master switch. Proceedings of the National Academy of Sciences. 2017, 114 (33) E6867–E6874; DOI:10.1073/pnas.1706915114

13. Zha Z, Bucher F, Nejatfard A, Zheng T, Zhang H, Yea K et al. Interferon-γ is a master checkpoint regulator of cytokine-induced differentiation. Proceedings of the National Academy of Sciences. 2017;114(33):E6867–E6874.

14. Deshmane S, Kremlev S, Amini S, Sawaya B. Monocyte Chemoattractant Protein-1 (MCP-1): An Overview. Journal of Interferon & Cytokine Research. 2009;29(6):313–326.

15. Sager R., Haskill S., Anisowicz A., Trask D., Pike M.C. (1991) GRO: A Novel Chemotactic Cytokine. In: Westwick J., Lindley I.J.D., Kunkel S.L. (eds) Chemotactic Cytokines. Advances in Experimental Medicine and Biology, vol 305. Springer, Boston, MA. https://doi.org/10.1007/978-1-4684-6009-4_9

16. Conti P, DiGioacchino M. MCP-1 and RANTES are mediators of acute and chronic inflammation. Allergy Asthma Proc. 2001 May-Jun;22(3):133–7. doi: 10.2500/108854101778148737. PMID: 11424873.

17. Knight BE, Kozlowski N, Havelin J, King T, Crocker SJ, Young EE and Baumbauer KM (2019) TIMP-1 Attenuates the Development of Inflammatory Pain Through MMP-Dependent and Receptor-Mediated Cell Signaling Mechanisms. Front. Mol. Neurosci. 12:220. doi: 10.3389/fnmol.2019.00220

18. Brindle NP, Saharinen P, Alitalo K. Signaling and functions of angiopoietin-1 in vascular protection. Circ Res. 2006;98(8):1014–1023. doi:10.1161/01.RES.0000218275.54089.12

19. Saharinen P, Alitalo K. The yin, the yang, and the Angiopoietin-1. Journal of Clinical Investigation. 2011;121(6):2157–2159.

20. Rybakowski J. Matrix Metalloproteinase-9 (MMP9)—A Mediating Enzyme in Cardiovascular Disease, Cancer, and Neuropsychiatric Disorders. Cardiovascular Psychiatry and Neurology. 2009;2009:1–7.

21. Nagase H, Visse R, Murphy G. Structure and function of matrix metalloproteinases and TIMPs. Cardiovascular Research. 2006;69(3):562–573.

22. Velazquez-Salinas L, Verdugo-Rodriguez A, Rodriguez LL and Borca MV (2019) The Role of Interleukin 6 During Viral Infections. Front. Microbiol. 10:1057. doi: 10.3389/fmicb.2019.01057

23. Tanaka T, Narazaki M, Kishimoto T. IL-6 in inflammation, immunity, and disease. Cold Spring Harb Perspect Biol. 2014;6(10):a016295. Published 2014 Sep 4. doi:10.1101/cshperspect.a016295

24. Kamburova EG, van den Hoogen MW, Koenen HJ, Baas MC, Hilbrands LB, Joosten I. Cytokine Release After Treatment With Rituximab in Renal Transplant Recipients. Transplantation. 2015 Sep;99(9):1907–11. doi: 10.1097/TP.0000000000000515. PMID: 25675201.

25. Couper K, Blount D, Riley E. IL-10: The Master Regulator of Immunity to Infection. The Journal of Immunology. 2008;180(9):5771–5777.

26. Di Paolo NC, Shayakhmetov DM. Interleukin 1α and the inflammatory process. Nat Immunol. 2016;17(8):906–913. doi:10.1038/ni.3503

27. Scholz A, Plate K, Reiss Y. Angiopoietin-2: a multifaceted cytokine that functions in both angiogenesis and inflammation. Annals of the New York Academy of Sciences. 2015;1347(1):45–51.

28. Shibuya M. Vascular endothelial growth factor (VEGF)-Receptor2: its biological functions, major signaling pathway, and specific ligand VEGF-E. Endothelium. 2006 Mar-Apr;13(2):63–9. doi: 10.1080/10623320600697955. PMID: 16728325.

29. Franzke A. The role of G-CSF in adaptive immunity. Cytokine Growth Factor Rev. 2006 Aug;17(4):235–44. doi: 10.1016/j.cytogfr.2006.05.002. Epub 2006 Jun 27. PMID: 16807060.

30. Weimin Wang, Dan Li, Liangliang Xiang, Mengying Lv, Li Tao, Tengyang Ni, Jianliang Deng, Xiancheng Gu, Sunagawa Masatara, Yanqing Liu & Yan Zhou (2019) TIMP-2 inhibits metastasis and predicts prognosis of colorectal cancer via regulating MMP-9. Cell Adhesion & Migration, 13:1, 272–283, DOI: 10.1080/19336918.2019.1639303

31. Ngu J, Meijndert C, Mewhort H, Turnbull J, Yong W, Stetler-Stevenson W et al. The Role of Tissue Inhibitor of Metalloproteinase-2 in Human Cardiac Fibroblast-Mediated Extracellular Matrix Remodeling. Canadian Journal of Cardiology. 2013;29(10):S363–S364.

32. Poniatowski L, Wojdasiewicz P, Gasik R, Szukiewicz D. Transforming Growth Factor Beta Family: Insight into the Role of Growth Factors in Regulation of Fracture Healing Biology and Potential Clinical Applications. Mediators of Inflammation. 2015;2015:1–17.

33. Morikawa M, Derynck R, Miyazono K. TGF-β and the TGF-β Family: Context-Dependent Roles in Cell and Tissue Physiology. Cold Spring Harbor Perspectives in Biology. 2016;8(5):a021873.

34. Bretscher PA. The activation and inactivation of mature CD4 T cells: a case for peripheral self-non self-discrimination. Scand J Immunol. 2014;79(6):348–360. doi:10.1111/sji.12173.

